# DEVELOPMENT OF A SOYBEAN MATURITY PREDICTION MODEL FOR SOYBEAN GROWN IN AFRICAN ENVIRONMENTS

**DOI:** 10.1101/2021.03.09.434647

**Authors:** Guillermo S. Marcillo, Nicolas F. Martin, Brian Diers, Michelle S. Da Fonseca, Erica Leles

## Abstract

Time-to-maturity (TTM) is an important trait in soybean breeding programs. However, soybean is a relatively new crop in Africa. As such, TTM information is not yet well defined as in other major producing areas. Multi Environment trials (MET) allow breeders to analyze crop performance across diverse conditions but also pose statistical challenges (e.g. unbalanced data). Modern statistical methods, e.g.. Generalized Additive Models (GAM), can flexibly smooth a range of responses while retaining observations that could be lost under other approaches. We leveraged 5 years of data from a MET breeding program in Africa to identify the best geographical and seasonal variables to explain site and genotypic differences in soybean TTM. Using soybean-cycle features (minimum temperature, daylength) along with trial geolocation (longitude, latitude), a GAM model predicted soybean TTM within ± 10 days of the average observed TTM [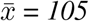 *days post-planting*]. Further, we found significant differences between cultivars (p<0.05) in TTM sensitivity to minimum temperature and daylength. Our results show promise to advance the design of maturity systems that enhance soybean planting and breeding decisions in Africa.

## 1 Introduction

Soybean (*Glycine max* (L.) Merr.) is a beneficial crop for smallholder agricultural systems in Africa. As part of a rotation, soybean can break yield limiting pathogen cycles (Carsky et al. 2000), and fix atmospheric N2-Nitrogen to reduce fertilizer requirements of subsequent grain crops. (Sinclair et al. 2014). Likewise, soybean stands out among legume species due the rich protein and oil contents of its seed. Soybean grown as a primary crop at a large scale would create options for enhanced security given its wide range of applications in the food and feed industry. From its early introduction in the 19th century, soybean planting area has increased from less than to 20,000 to 1,500,000 ha. by the late 2010’s (Khojely et al. 2018). This expansion has occurred presumably due to the significant value of the crop in regional trade networks, which strengthens domestic production, reduce the demand for imports, and even favor surpluses for exports (Foyer et al. 2019; Keyser and Van Gent 2007). Despite the potential for increases in soybean production, research is still needed to address productivity challenges that African growers face to date, such as declined soil fertility, poor farm practices, and low-yielding cultivars.

The Soybean Innovation Lab (SIL) (“Soybean Innovation Lab” 2020) is a USAID funded program focused on advancing soybean in Africa. The Pan African Variety Trials (SIL-PAT) (“Tropical Soybean Information Portal” 2020) is a multienvionment soybean trial network currently conducting trials over 100 locations in 24 countries. SIL-PAT partners with public and private organizations to test commercial soybean cultivars sourced from across Africa, the US, Australia, and Latin-America (Santos 2019). To date, trials carried out at SIL-PAT have enabled the registration of 7 new soybean varieties in Ghana, Ethiopia, Malawi, Mali, and Uganda, and 10 more are in the process for Cameroon, Ethiopia, Kenya, Malawi, and Zambia (Leles 2021). Aside from yield, SIL-PAT also hosts a database with volumes of phenotypic observations related to agronomic and nutritional quality. The SIL-PAT database offers a unique opportunity to collate disperse multi-environmental trial datasets, which can enable the characterization of soybean performance across diverse cropping conditions in the Pan-African region. Among these traits, the time-to-maturity (TTM) of a soybean cultivar is directly related to the commercial cycle’s length expected for new cultivars introduced to the market.

Time to Maturity (TTM) is associated with the biological length cycle of a cultivar (Ersoz, Martin, and Stapleton 2020). Therefore, increasing the understanding of the factors that influence it is critical to define the geographical adaptation for new cultivars. More specifically, the expected TTM of a cultivar will depend on the conditions prevalent during the growing cycle such as daylength and temperature. In hemispheric areas, for example, soybean maturity is delayed from lower to higher latitude locations. Thus, cultivars adapted to southern latitudes are expected to respond better to shorter days than cultivars adapted to the northern region (Zhang et al. 2007; Mourtzinis and Conley 2017). In addition, temperature has been reported to influence TTM and post-vegetative soybean development in general ; with studies documenting early flowering occurring under higher spring temperatures (Cooper 2003), or pre and post-flowering development rates being affected by the interaction of photoperiod, temperature, and genotype (Cober, Stewart, and Voldeng 2001). Acknowledging daylength and temperature as the prime drivers of reproductive development has been important to delineate areas for soybean adaptation in Northern latitudes (Scott and Aldrich 1983). Furthermore, a careful distinction of daylength and temperature effects has permitted to identify optimal areas of adaptation, with a consequent impact on resource allocation and soybean productivity in the Northern and Southern latitudes. On the contrary, field evidence of thermal and photoperiodic effects on the onset of physiological maturity is rather limited for emerging soybean markets in the tropics, such as the Pan-African region. Moreover, no previous characterizations of TTM have been released over a large geographical coverage in Africa.

Using 5-year data (2015-2020) from 150 cultivars and experimental lines evaluated at 65 sites in the SIL PAT network, we set the following goals : 1) Identify the best combination of geographical and seasonal characterization variables (i.e. elevation, latitude, longitude, temperature, and daylength) that explained site and genotypic differences in soybean TTM; and, 2) Evaluate the usability of these variables to build a parsimonious predictive model of soybean maturity timing adapted to the growing conditions in the Pan-African region. Results from this research will be used to understand cultivar by environment interactions as well as support the selection of cultivars adapted to African farmer fields. Our work will lay ground for building a maturity classification system, currently missing for soybean growers in Africa. Knowing maturity timing in advance is important for growers to improve their planting decisions, and for breeders to best plan their trials.

## 2 Materials and Methods

### 2.1 Maturity time explained by genotype and environment

In a first step we performed exploratory analysis of soybean maturity times. Sequential three-way ANOVA models were used to evaluate the main sources of variability in time to Maturity TTM (days after planting) due to the additive effects of genotype (G), Location (L), Season (S), and the combined effects of location (L) and season (S) into the factor Environment (E). Next, mean estimates for TTM were obtained by adjusting random effects for G and E. Observations which departed unusually from linear model assumptions were identified and excluded on the basis of influential observations (Cook’s distance analysis) and residuals vs. fitted maturity-time plots. Mixed effects models have proven effective to analyze phenotypic data generated in a breeding program (Bernardo 2020). Best Linear Unbiased Predictions (BLUPS) from this process were used as the target variable to predict soybean TTM in response to explicit environmental queues in a second step. The two-stage analysis was computationally efficient. It described adequately within-environment errors first (Piepho et al. 2012) and then focused on prediction accuracy in the second stage. Stagewise approaches allow for adjusting cultivar means per trial for later analyses and enable combined analyses of large amounts data carrying significant variation across environments (Buntaran et al. 2020).

### 2.2 Modeling Maturity as a function of environment

Weather records were spatially linked to the geographic coordinates of the trialing sites in SIL PAT. Soybean cropping conditions were characterized by the geographic and meteorological variables recorded during the length cycle. Temperature and daylength are the most physiologically meaningful drivers of phenological changes in soybean (Major et al. 1975) and were included in this analysis as such. Daily meteorological variables were averaged or summed from planting up to the occurrence of three phenological stages, i.e. emergence, flowering initiation, and physiological Maturity. Minimum, maximum, and mean daily temperatures, [*°*C] were provided by aWhere (“AWhere | Climate Smart Weather Insights Backed by AI” 2021) and validated to ancillary station-based data. Daylength [hours. day^−^1] was simulated as a function of latitude based off standard equations by Campbell and Norman (Campbell and Norman 2000) and Teh (Teh 2006). Aside from absolute values for temperatures and daylength, we considered additional variables capturing the differences in maximum, minimum, and mean daily temperature, and daylength, computed in between-stages. For example, DLMEANDIFF signifies the difference in hours of light received from flowering to Maturity for a certain cultivar at a given location. A positive value for DLMEANDIFF is the result of more hours of light received at flowering than Maturity and implies that a soybean cultivar planted at a certain location was growing as days were getting shorter. In contrast, a negative value balances the difference in favor of the daylength received at Maturity, and thus the soybean cycle of a given cultivar progressed along longer days. It must be noted that the SIL PAT trials were conducted within a considerable latitudinal range (i.e. [-21*°*S, 13*°*N]), but still circumscribed to the tropics. As such, the post-flowering daylength period at the trialing sites varied between ([-0.51, +0.72], hours). A full description of the variables considered for soybean TTM prediction is presented in supplementary materials.

As a next step, we ran forward stepwise regression to identify the most essential variables that could be combined in a predictive model of soybean TTM. Redundancy in this set was reduced by removing highly collinear variables via Spearman ranking correlation. Different feature sets combining a temperature-based predictor at the time along with DLMEANDIFF and geolocation (lat, long) were evaluated independently. The ‘Mallow’s (Cp), ‘Hocking’s (Sp), and ‘Amemiya’s prediction (APC) indexes (Amemiya 1980) were used to select the best feature combination. Both Cp and Sp measure the fraction of variability in the response variable (i.e. residual sum of squares, RSS) that results from recursively fitting models with one regressor removed at a time. The APC index is an adjusted R^2^ that penalizes additional parameters (i.e. degrees of freedom) in the regression’s right-hand side. Lower values for Cp and Sp, and higher values for APC, are equivalent and indicative of a better model. Variables in the final subset were used as predictors for fitting and parametrizing a Generalized Additive Model (GAM) to predict soybean TTM. Traditional breeding techniques are usually performed on reduced MET datasets due to the few genotypes that get retained for evaluation in late stages of the trialing process (Dawson et al. 2013). In our analysis, we avoided losing indiscriminately far too many observations and favored the application of the GAM statistical algorithm to capture data signals that could be lost with the use of alternative approaches.

The GAM algorithm (James et al. 2013) has been traditionally applied to problems in ecology, land allocation, and climatology (Roberts and Key 2008; Stauffer et al. 2017; Lawler et al. 2006). Given its flexibility and simplicity to capture complex responses, it is being implemented more often to predict field-level agricultural traits, for example wheat yield (Chen, O’Leary, and Evans 2019), pasture biomass (De Rosa et al. 2021), or pest use assement (Rosenheim et al. 2020). Our study is the first documented application of a GAM model to predict soybean phenotypic traits across tropical environments. A general specification of the final GAM maturity model followed:

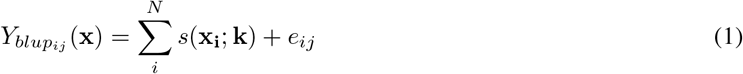

The response variable Y_*blup*_ in equation 1 was the mean TTM previously adjusted for genotype, location, and season effects. Y_*blup*_ is predicted with an additive function of the best environmental features (e.g. mean temperature to Maturity, daylength, etc.). Each predictor (x) is smoothed by a basis function *s(*.) that modulates the maturity response through parameter k. The k-parameter is a knot indicating whether there is a change in the direction of the response and could be tuned as in other non-parametric approaches such as splines. The GAM model’s advantage is that non-linear relationships carried by the predictors can be easily smoothed to improve model fit without increasing complexity as in other parametric approaches (e.g. nonlinear multivariate regression). Model parametrization was sequential and used a 5-fold cross validation with 5 replicates to find the optimal knot (k-parameter) for each predictor at a time. For model training and validation, we balanced the number of observations in each dataset ensuring that trial planting-dates be sufficiently represented. In this fashion, the training dataset included observations for 2018, 2019 (summer seasons), and 2018-2019, 2019-2020 (winter seasons). Seasons 2016-2017 and 2017-2018 were held-out for model validation. To account for spatial variability, latitude and longitude coordinates of each trial were also included as predictors.

## 3 Results and Discussion

### 3.1 Exploratory analysis of soybean Maturity timing

A sample size of 2,823 observations (Genotype x Location x Season) was used to analyze the sources of variability of the TTM trait recorded in the SIL-PAT dataset (Figure 1, Figure 2). Genotype (G) and Location (L) separately explained 12 or 68% of the total variability in maturity time. While the contribution of cropping Season (S) alone was low, it helped to account for almost 87% of differences in maturity timing across genotypes and locations. Further, environmental effects (Location + Season) were almost six times those of genotype as evidenced by adjusted R^2^, and type II sum of squares (SS) estimated in sequential ANOVA models fitted to time-to-maturity. Expectedly, there was also a gradual decrease in the standard error for the TTM residuals (RSE, Table 1) as additional sources of variability were considered.

**Table 1:** Soybean time to maturity TTM, sources of variation

| Factor | <sup>2</sup>DF Model | DF Residuals | <sup>3</sup>RSS | Adj-R<sup>2</sup> | <sup>4</sup>RSE (days) |
| --- | --- | --- | --- | --- | --- |
| Genotype (G) | 174 | 2,648 | 882,623 | 0.12 | 18.2 |
| Location (L) | 67 | 2,755 | 330,979 | 0.68 | 10.9 |
| Season (S) | 8 | 2,814 | 1,017,289 | 0.05 | 19 |
| G + L | 241 | 2,581 | 225,891 | 0.76 | 9.3 |
| G + S | 182 | 2,640 | 864,933 | 0.14 | 18.1 |
| E= L + S | 75 | 2,747 | 280,997 | 0.73 | 10.1 |
| G + E | 265 | 2,557 | 121,887 | 0.87 | 6.9 |

<sup>2</sup> DF= Degrees of freedom, <sup>3</sup>RSS= Type II Residual sum of squares, <sup>4</sup> RSE= Residuals standard error

**Figure 1:**
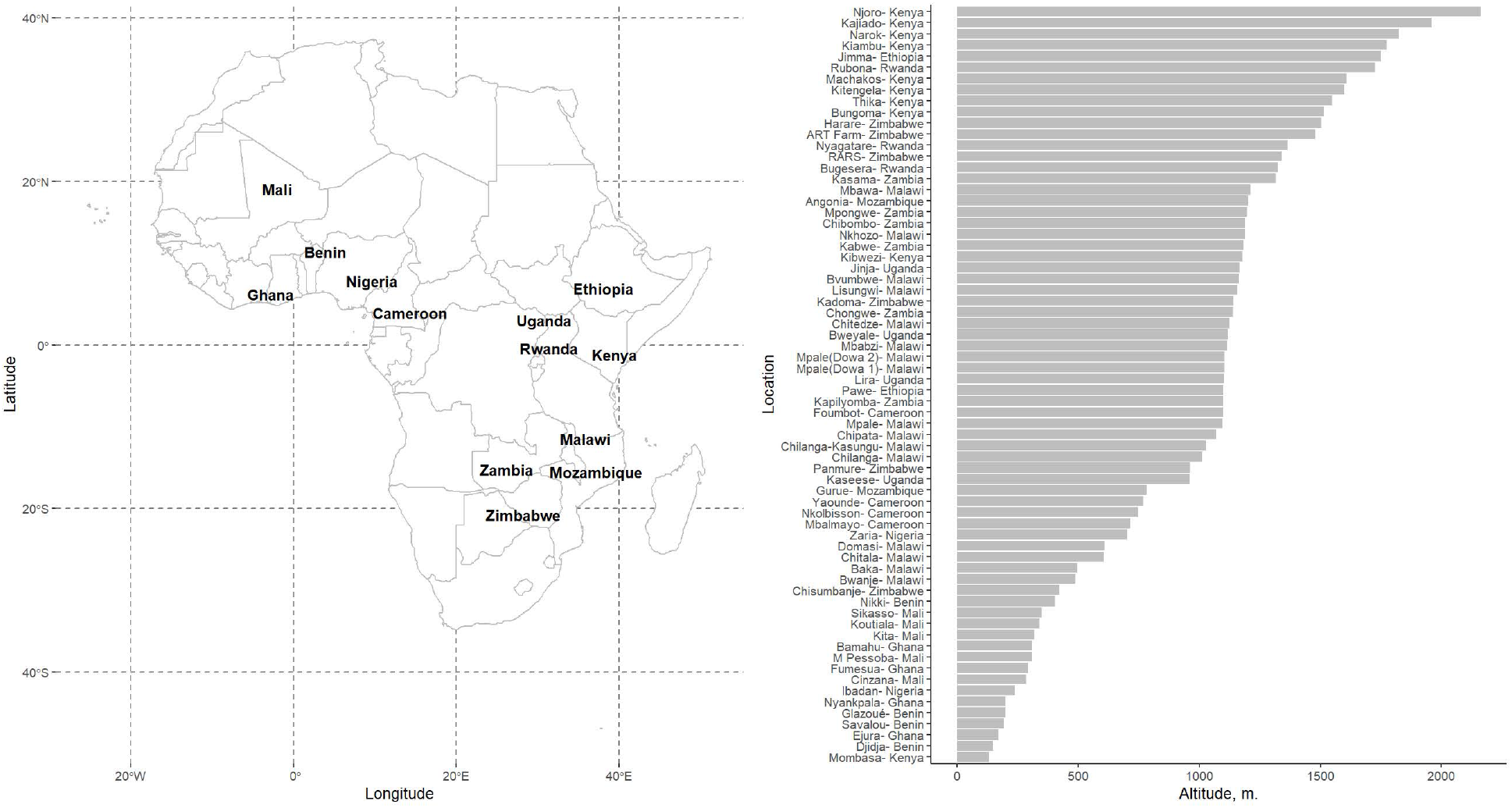
Countries and trial site allocation in the Soybean Innovation Lab, Pan-African trials (SIL-PAT, 2015-2020)

**Figure 2:**
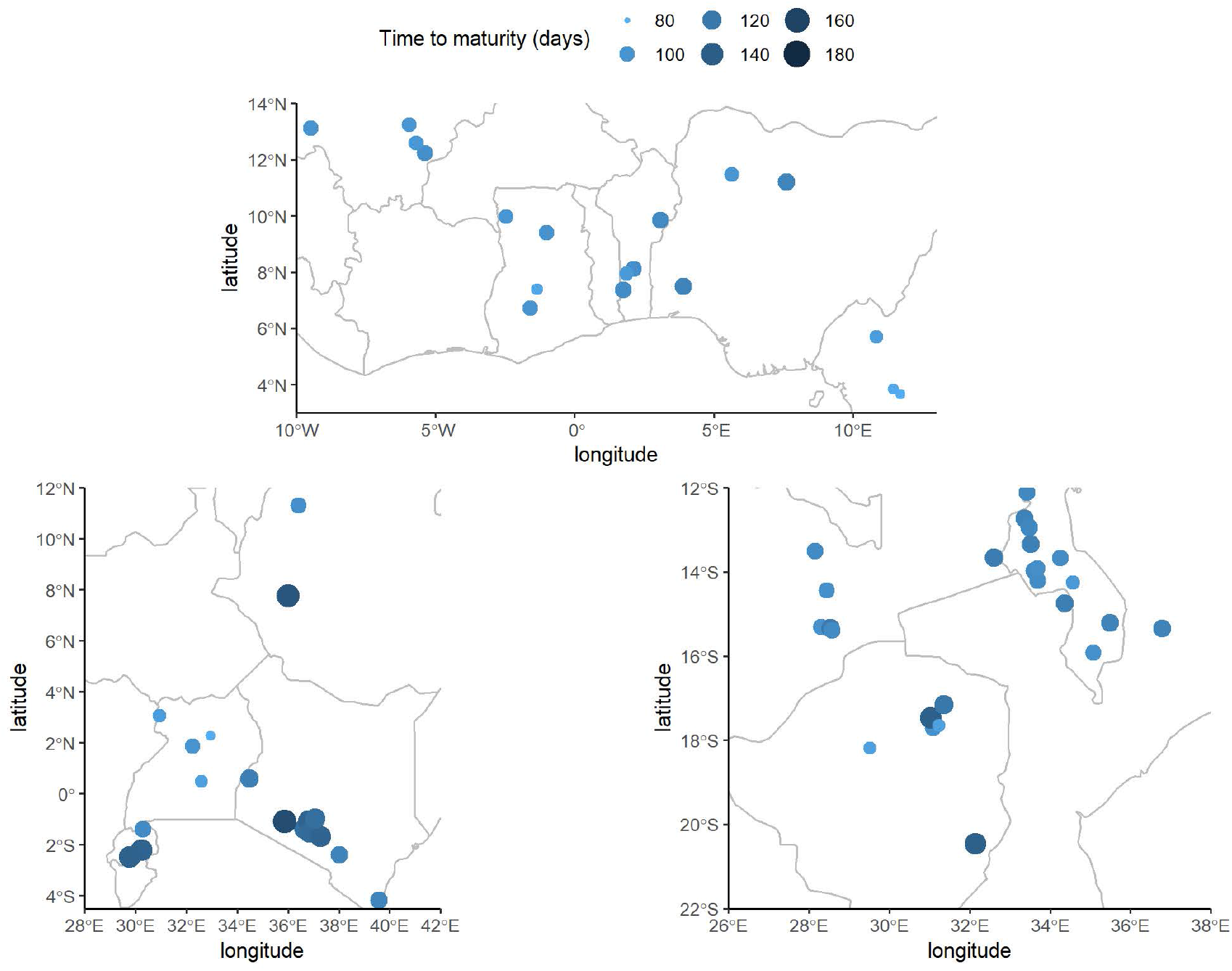
Geographical variation of soybean time to maturity in the Pan-African variety trial network (SIL-PAT).

Mean TTM was adjusted for hierarchies in the SIL PAT dataset by means of random-effects modeling. The random-effects model captured discrepancies efficiently as 98.5% of the resulting time-to-maturity BLUPS met model assumptions (i.e. residuals randomly scattered and bounded within three times the residual SE, Figure 3). Additional details on outlier detection through residual and Cook’s distance analysis can be found in the supplementary materials. The outliers removed corresponded to the L342-Chilanga-2019, N390-Chilanga-2019, and SP 8 DPSB – Thika 2016/2017 genotype by environment entries. The overall mean TTM was 108.9 days after planting [95% CI: 105-113 days]. Around the estimated mean, time-to-maturity departed by 8 or 19 days due to G and E effects, respectively (Table 2). The significant pool of variation in maturity occurrence across genotypes and locations in SIL PAT (i.e. 89%, Table 2) warranted the exploration of environmental queues that can be used in a parsimonious model to predict maturity times in Sub-Saharan Africa.

**Table 2:**
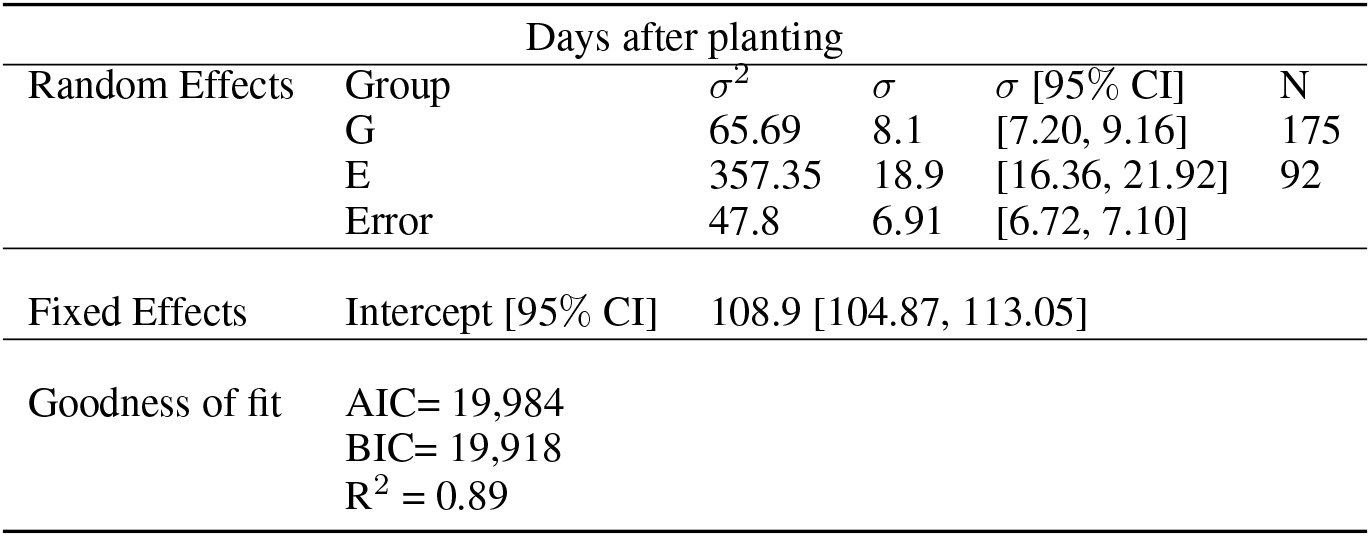
Soybean time to maturity adjusted by genotype (G) and environment (E) effects

**Figure 3:**
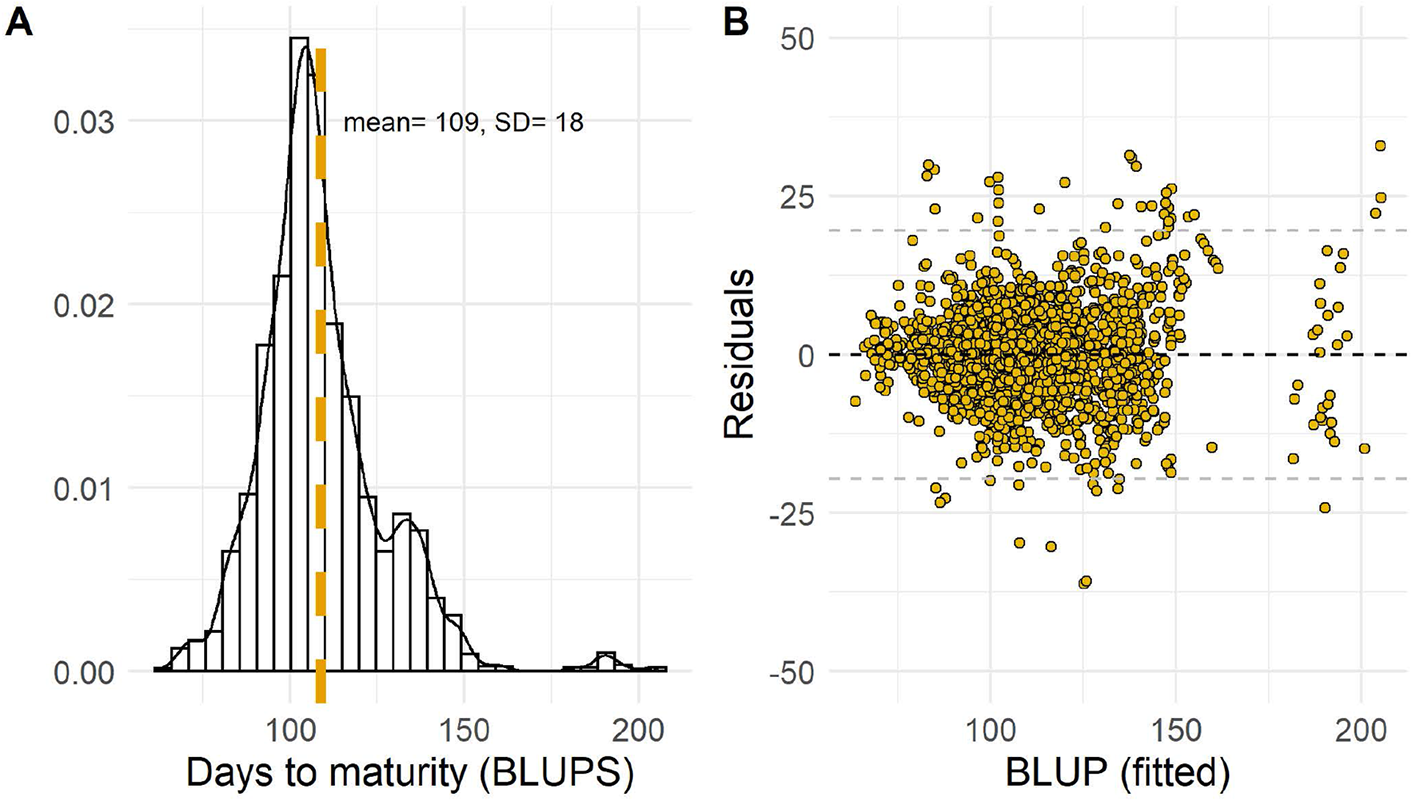
Time to maturity TTM, Best Linear Unbiased predictions. Panel (B) displays residuals of the random-effects model. Horizontal dashed lines represent upper and lower threshold limits bounded within ± 3 SE

### 3.2 Best features to characterize soybean time to maturity TTM

The best features to explain changes in TTM were the daily minimum temperature from planting to Maturity (TMINM), and the difference in daylength from flowering to Maturity (DLMEANDIFF). Accounting also for the effects of latitude and longitude, the best subset captured 36% of the differences in Maturity reported across genotypes and locations in the SIL PAT dataset (Table 3). The actual and fitted responses of time to Maturity to each predictor are visualized in figure 4. Likewise, the best feature subset displayed the lowest numbers for AIC, Cp, HSP, and AP, suggesting that complexity (i.e. number of parameters) and explanatory capabilities were balanced in the GAM model to predict soybean TTM that was built on the basis of these variables. The GAM model using the best explanatories of soybean improved the accuracy of TTM predictions (Table 4). The GAM model used three “break points” (i.e. k-nodes) to smooth the overall negative relationship between TTM and each of TMINM and DLMEANDIFF. While latitude and longitude show less than a strong association with the response (figure 4), including a two-dimensional smooth for these terms helped account for the spatial variability due to trial location (Table 4). Model fit improved as a result.

**Table 3:**
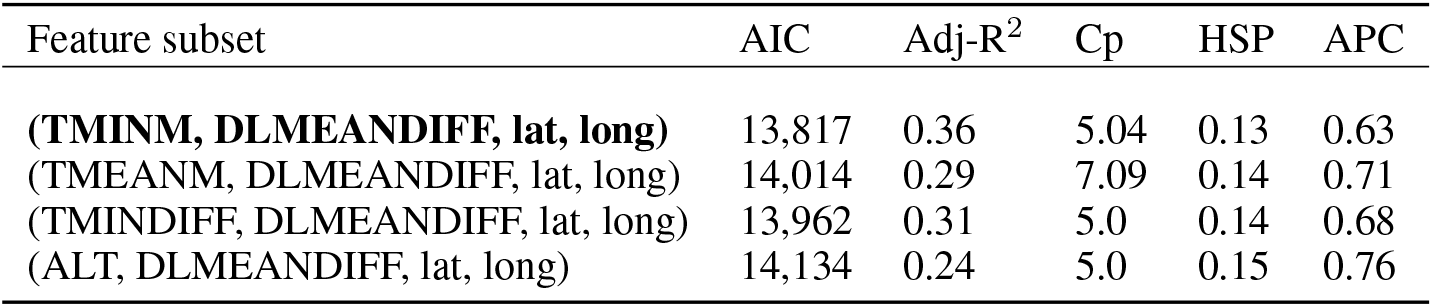
Best subset of features to be used as predictors in a soybean time-to-maturity model.

**Table 4:** Evaluation and testing of GAM models used to predict Soybean maturity timing. Linear regression (lm) models disaplyed for comparison

| Model | 5-fold Validation |  |  |  | AIC | BIC |
| --- | --- | --- | --- | --- | --- | --- |
|  | RMSE (days) |  | R<sup>2</sup> |  |  |  |
|  | Training | Testing | Training | Testing |  |  |
| lm ~ TMINM | 15.34 | 15.79 | 0.29 | 0.30 | 14,000 | 14,016 |
| gam ~ s(TMINM, 3) | 13.55 | 13.67 | 0.46 | 0.46 | 13,542 | 13,564 |
| lm ~ s(TMINM, DLMDIFF) | 14.96 | 15.48 | 0.32 | 0.33 | 13,924 | 13,945 |
| gam ~ s(TMINM, 3) + DLMDIFF | 12.50 | 12.70 | 0.54 | 0.53 | 13,280 | 13,307 |
| gam ~ s(TMINM, 3) + s(DLMDIFF, 3) | 10.78 | 11.02 | 0.66 | 0.65 | 12,793 | 12,825 |
| lm ~ (TMINM + DLMDIFF + lat + long) | 14.90 | 15.48 | 0.34 | 0.32 | 13,817 | 13,850 |
| gam ~ s(TMINM, 3) + s(DLMDIFF, 3) + lat + long | 10.05 | 10.3 | 0.70 | 0.69 | 12,576 | 12,620 |
| **gam ~ s(TMINM, 3) + s(DLMDIFF, 3) + s(lat, long, 4)** | **9.99** | **10.35** | **0.70** | **0.69** | **12,564** | **12,613** |

**Figure 4:**
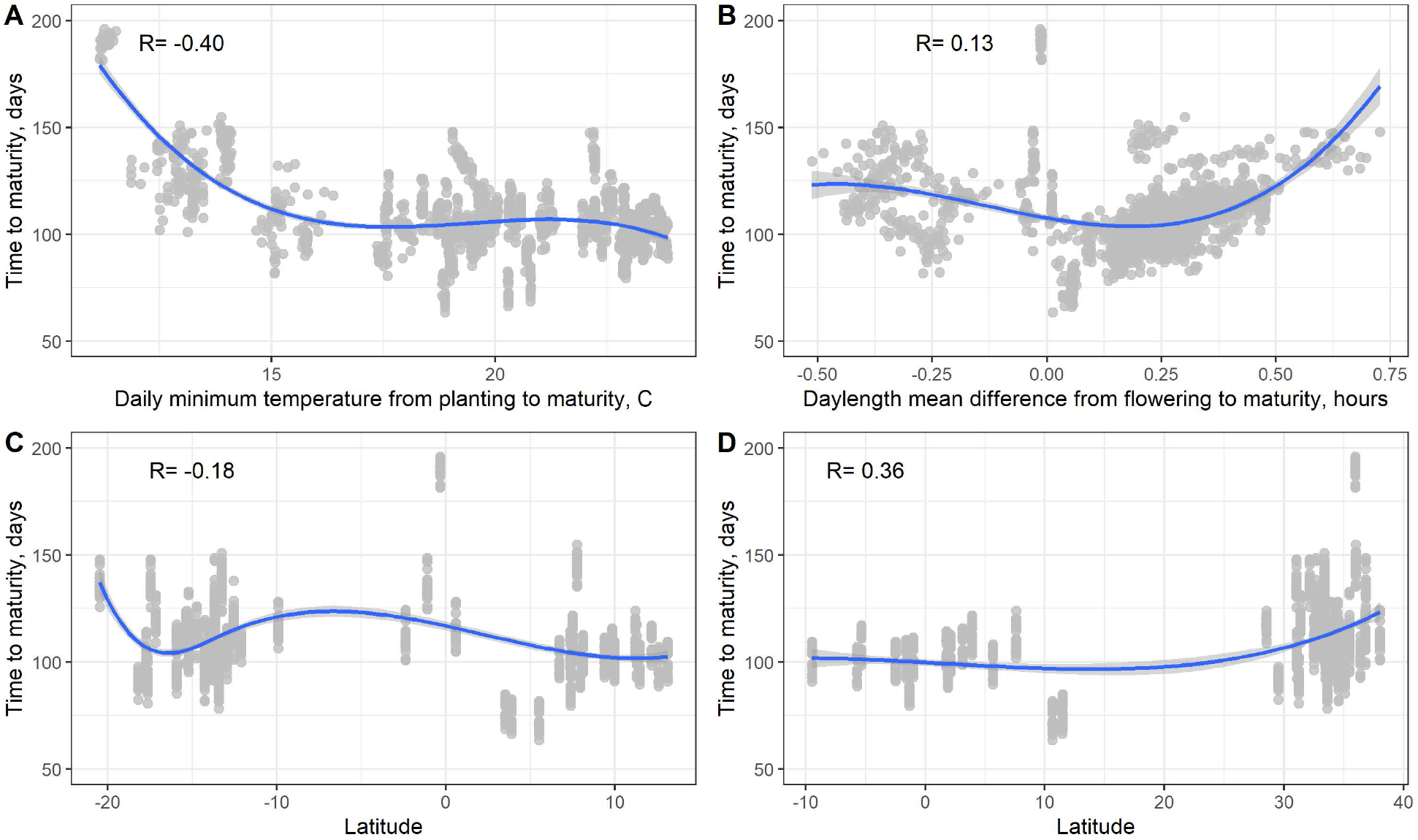
Time to maturity response (BLUP) to weather and location variables used as the best predictors in the GAM model. Non-linear effects for each variable visualized with natural splines functions with a knot number equal to three.

Following 5-fold model validation, the overall expected TTM for a cultivar tested in the SIL PAT network was predicted within ±10 days of the observed field data. Relative to simple linear regression, prediction error with the GAM model decreased by almost 33%. Likewise, R^2^ increased to nearly 70% whereas AIC was the lowest among several specifications of the GAM model. A more detailed description model-agreement during the training and testing phases is presented in Figure 5 and all the models considered can be found in supplementary materials. GAM predictions fitted acceptably well with the observed time to Maturity (Figure 5). RMSE ranged between 9 and 14 days at different soybean growing seasons considered in the validation sets. Relative to other seasons, the model under and over predicted time-to-maturity in 2016/2017 and 2018 by 4% relative to the observed overall mean 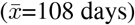. The lower fit in 2018 was associated with cultivars tested in mid to high elevation sites in Rwanda ([1,300-1,700] m). In contrast, the less than ideal fit in 2016/2017 was presumably due to fewer cultivars tested at very low elevations in Mali ([280-320] m). Overall, soybean TTM predictions held acceptably well for 70% of the 122 genotypes considered for training and testing the GAM model (±5 days off the observed Maturity). The remaining 30% of the cultivars were off by 6 days or more and resulted from low sampling, i.e. 15 sites evaluated or fewer.

**Figure 5:**
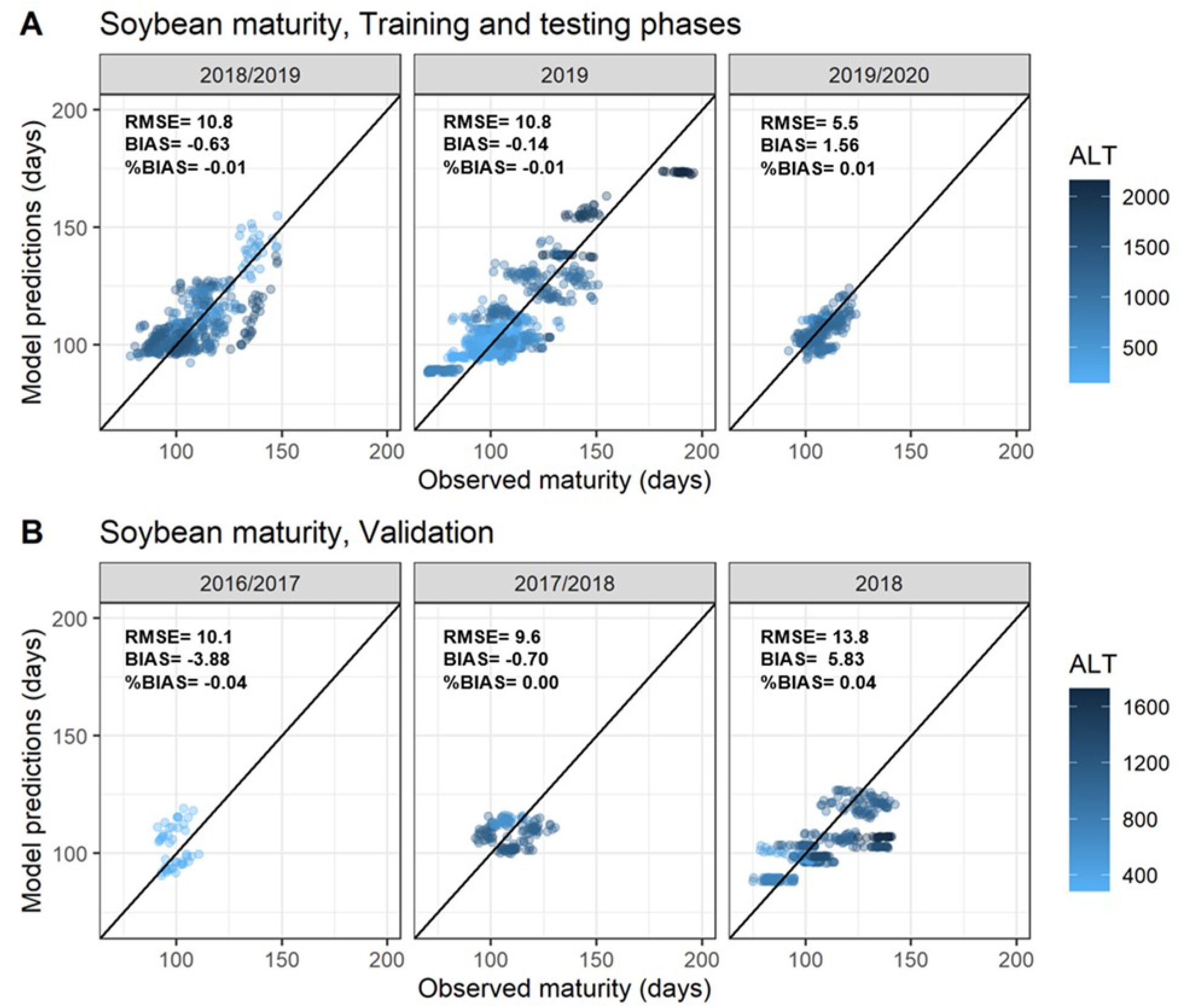
GAM Model agreement during the training and validation phases. The color bar on the right indicates the range of elevations (ALT, m).

### 3.3 Soybean maturity responses to temperature and daylength in Africa

Cultivars that were tested more consistently across environments (n= 33) captured a larger and more significant range of TTM response to minimum temperature (TMINM) and daylength (DLMEANDIFF). These cultivars were also part of foreign germplasm introduced with potential for fast introduction to the African markets. Critical points where a shift in TTM occurred in response to TMINM and DLMEANDIFF were approximated for each cultivar by means of segmented regression (Muggeo 2008; Küchenhoff and Carroll 1997). These critical values delimit ranges of sensitivity for the TTM response to TMINM and DLMEANDIFF. To illustrate (Figure 6), the cultivar “TGX 2014-16FM” reached maturity around 105 days post-planting when tested in sites whose minimum temperatures during the cycle were between 17 and 30 °C. In turn, Maturity was sharply delayed by almost 60 days in sites within the 12-17 °C range. A similar analysis revealed that the cultivar “Lukanga” displayed a slightly higher value of thermal sensitivity (i.e. 19 °C). For “Lukanga”, seemingly late and early patterns of Maturity occurred shortly before 19 °C and thereafter.

**Figure 6:**
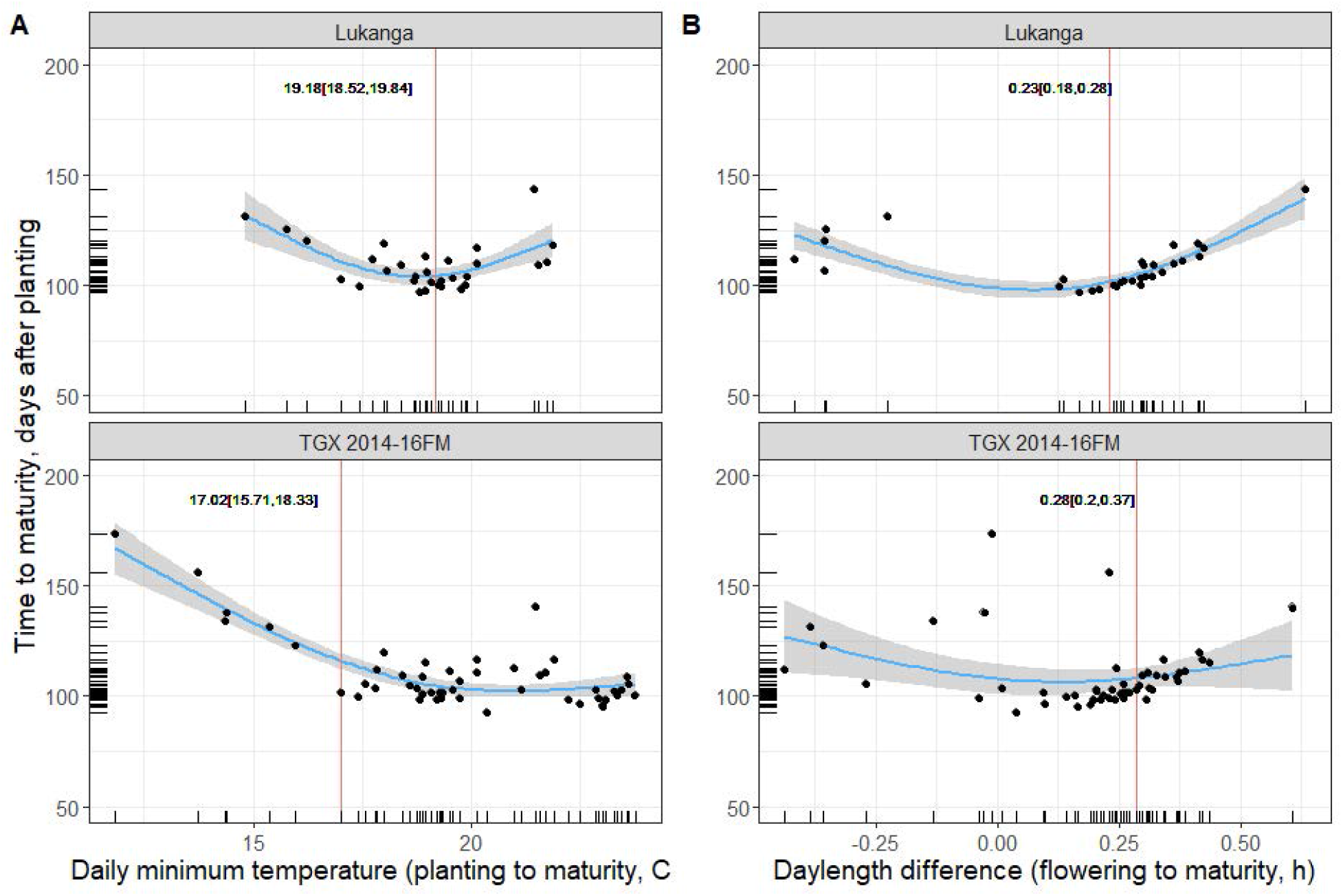
GAM smoothed response of soybean maturity time to environmental factors. Each dot represents a testing site where the cultivar was tested. Red parallel lines represent a change in the direction of the response, approximated using segmented regression.

In the same vein, physiological Maturity occurred more prevalently around 100 days post-planting when the daylength interval between Flowering and Maturity (DLMEANDIFF) approached +0.20 hours. A positive value for this explanatory variable means that the given cultivars received more light-hours at flowering than Maturity and is indicative of their growing cycles progressing along shorter days. As in the case of minimum temperature, cultivars seemed to follow different patterns of TTM response when the gap in daylength between flowering and Maturity moved far from a critical value. In the cultivars shown, the critical point of sensitivity to daylength were estimated at +0.23 and +0.28 hours of light. Based on an extended analysis, all the cultivars considered for analysis could be ranked in terms of their sensitivities to temperature and daylength as more trials are conducted in the coming years by the SIL-PAT network.

### 3.4 Discussion and implications

The environment and genotype effects were significant sources of variation in the time to maturity trait. Roughly, the environment (location x season) effects were almost six times the effects carried by genotype. Our findings are the first to systematically quantify location and genotype effects on soybean maturity time in Africa. Our results corroborate other reports from sub-tropical regions; areas characterized by short days and long summers. Alliprandini et al., (2009), for instance, found that location and genotype accounted for 62% and 29% of the variance in number of days to Maturity recorded for commercial cultivars adapted to recurrent cultivation ecoregions in Brazil (Alliprandini et al. 2009). While a significant main effect of environment is important in characterizing a phenotype response, between-cultivar differences are more informative from a breeding perspective (Malosetti, Ribaut, and van Eeuwijk 2013). Furthermore, genotype and genotype-by-environment (GE) interactions are ubiquitous in phenotypic characterization studies.

GE effects were revealed by cultivar specific patterns of Maturity that emerged from smoothing the responses attributed to weather features in the GAM model. Our results contribute to increasing the understanding of joint effects of temperature and daylength on the phasic development of soybean adapted to the African region. Reports from tropical conditions within the same latitudinal circumscription, such as Hawaii, showed that colder nights (i.e. minimum temperatures in high elevations) extended soybean vegetative periods and delayed physiological maturity (R7) by 25 days (George, Bartholomew, and Singleton 1990). More importantly, we highlighted the seemingly higher importance that thermal variation had relative to photoperiod in characterizing maturity in less hemispheric areas. In fact, low temperature effects on soybean field development unfold when photoperiodic effects exist but are minimum (Lawn and Byth 1974). A close inspection into critical values of response for these areas helped to define ranges of response where maturity of a cultivar would occur more or less consistently. Such ranges may indicate areas of geographical adaptation for a given cultivar. In turn, sudden shifts in maturity are associated with growing conditions in cultivation areas outside the possible ranges of adaptation. A cultivar planted under extreme conditions would mature either too early or too late. Consequently, the asynchronous occurrence of maturity leads to incomplete cycles with detriments on production. Soybean yields, for example, tend to be the highest for cultivars that maximize resources during the whole growth cycle (Egli 1993). Low yields can also be the result of premature maturity in short-stature plants that flower too early (Sinclair and Hinson 1992).

Soybean maturity time in the SIL-PAT network can be accurately predicted using geographical and seasonal characterization variables. Statistical learning can assist in the construction of parsimonious models that replace complex approaches, such as mechanistic models, to readily assist soybean field operations. Mechanistic, or process-oriented, models can be arguably more accurate under particular conditions but require a full detailed description of the physiological processes involved in plant development and growth (i.e. parametrization). Possible accuracy losses from a statistical model are compensated by the less demanding need of inputs and ease adaptation to other regions. Accordingly, models need to be constantly updated as the volume of information in their inputs increase. The expansion of the SIL PAT Network, and the information provided within it, will facilitate the validation of the findings from this study. Likewise, it is expected that model would extend the estimation of soybean TTM to locations in Africa not considered initially in this study.

## SUPPLEMENTARY MATERIALS

### S1. Analysis of residuals and influential observations

Cook’s coefficients are distance measures calculated as the residual obtained by fitting a model with all the observations included relative to one with an observation removed at the time. This difference is normalized by the inverse of an error or cost metric (e.g. RMSE). A higher value for the coefficient indicates a higher likelihood for an observation being too much influential in the overall estimation of model parameters. Distance metrics were adjusted for genotype-grouping level effects.

**S1 figure.**
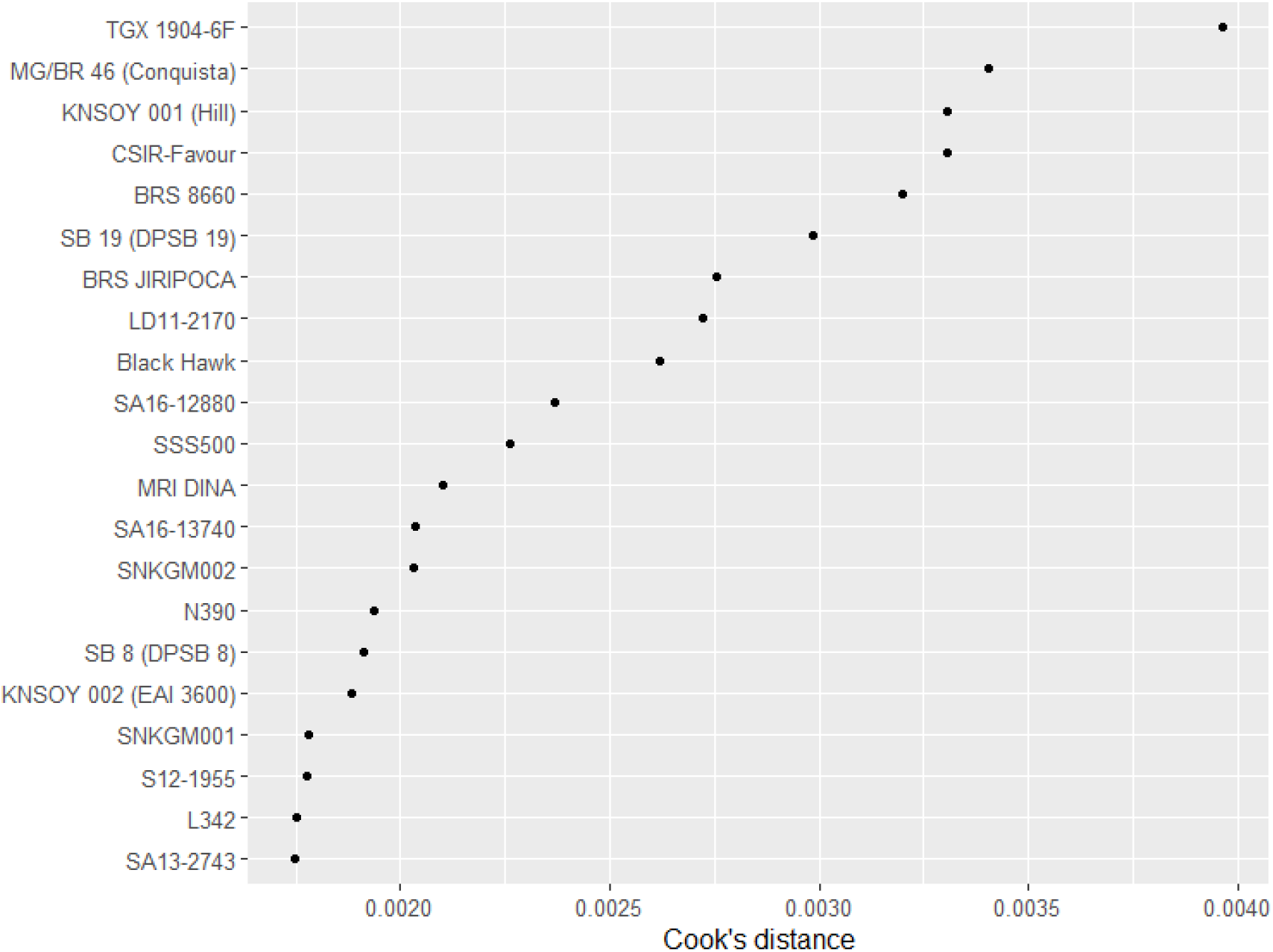
Ranking of top influential genotypes sorted by Cook’s coefficient values

Influential analysis based on Cook’s distance should not necessarily lead to automatic deletion of an observation. Here, we complemented the influence analysis with visual inspection of residuals. We assumed an acceptance threshold region bounded by 3-times the standard error of the residuals from the random effects model. Overall, 28 outliers (GxLxS) departed from this assumption (1.5 % of all BLUPS). Furthermore, only three out of 21 genotypes with a large distance coefficient were part of an environment with a large residual. (L342-Chilanga-2019, N390-Chilanga-2019, and SP 8 DPSB – Thika 2016/2017). The random effects model was reestimated with these three observations removed.

**S2 figure.**
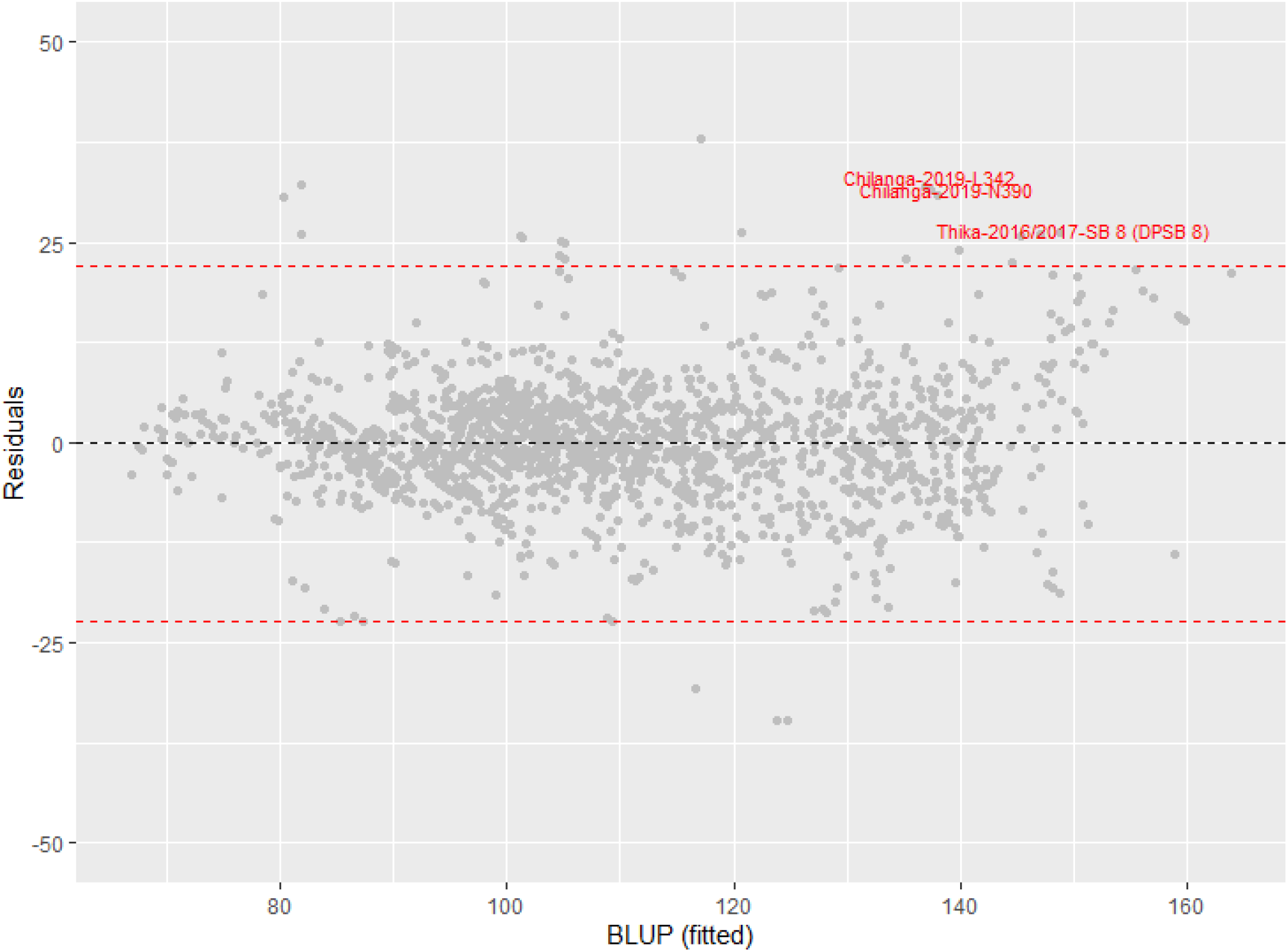
Outliers corresponding to observations displaying both a large Cook’s coefficient and a large residual.

### S2. Modeling: Training/testing sets based on planting date availability

**S3 figure.**
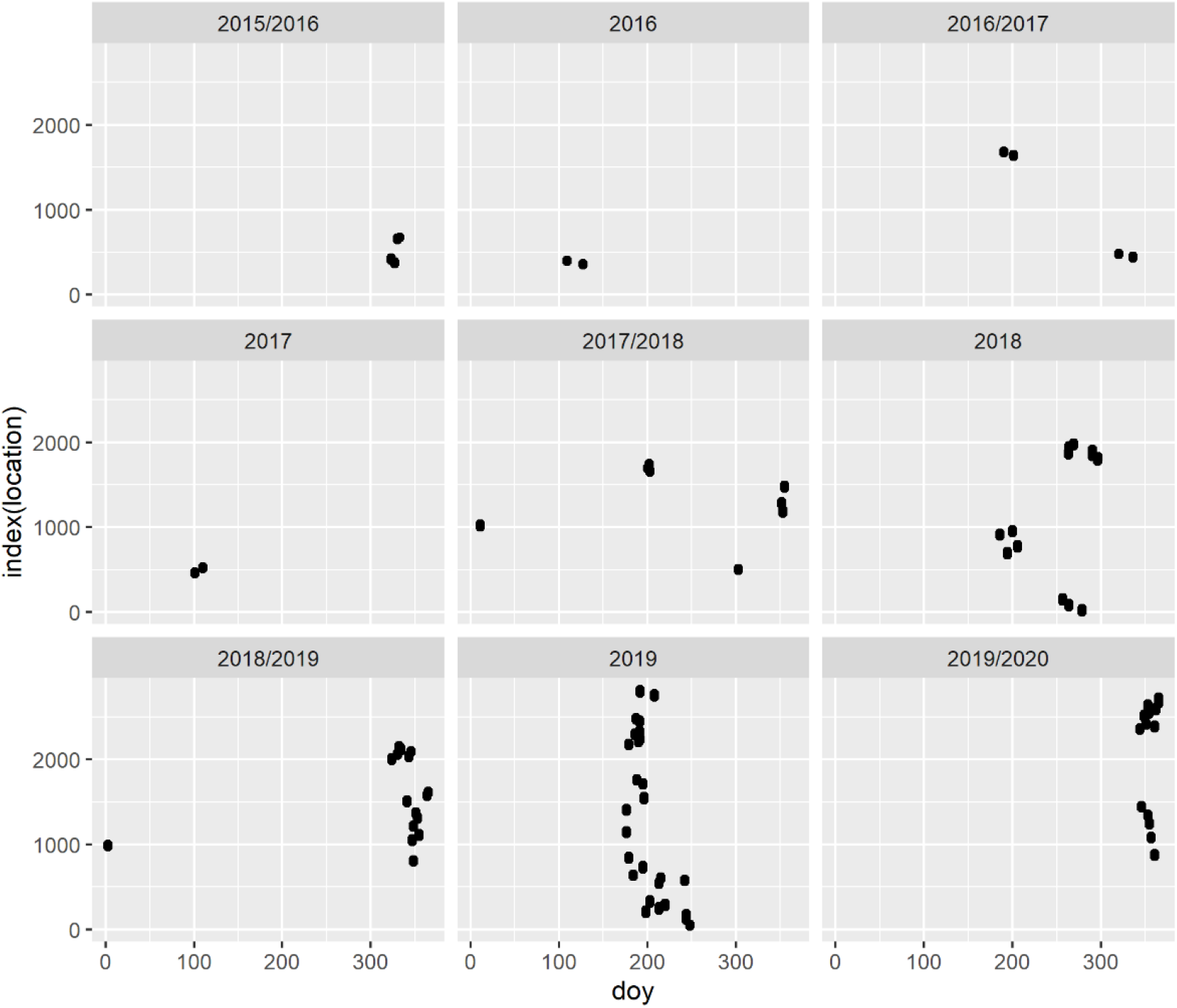
Planting date variation in the SIL -PAT trials across years. Planting dates converted to Julian-days (doy). Winter crops planted between-years (∼ day of year > 300). Summer crops planted around mid-months within a year (∼ day of year 200). To account for this variation and use the most information available during the training phase, the training and testing datasets included at least a winter and summer soybean crop for the 2018/2019, 2019, and 2019/2020 seasons. Data from previous years were held-out for model validation

### S2. Modelling: Features considered for Soybean TTM prediction

S2. Modeling: Features considered for Soybean TTM prediction
| Temperature (° C) |  |  |  |
| --- | --- | --- | --- |
| Label | Description | Mean (<sup>1</sup>SD) | Range (<sup>2</sup>IQR) |
| TMAXF | Daily maximum temperature to flowering | 28.1(2.0) | 22.4-32.5 (2.4) |
| TMINMF | Daily minimum temperature to flowering | 19.7 (3.2) | 11.2-23.9 (3.8) |
| TMEANF | Daily mean temperature to flowering | 23.9 (2.5) | 16.8-28.2 (3.3) |
| TMAXM | Daily maximum temperature to maturity | 28.5 (2.0) | 22.5-32.9 (2.8) |
| TMINMM | Daily minimum temperature to maturity | 19.8 (2.8) | 11.8-23.9 (3.8) |
| TMEANM | Daily mean temperature to maturity | 24.1 (2.3) | 17.2-28.4 (3.4) |
| Daylength (hours) |  |  |  |
| DLMEANF | Mean daylength to flowering | 12.5 (0.5) | 11.4-13.3 (0.5) |
| DLMEANM | Mean daylength to maturity | 12.4 (0.3) | 11.7-13.1 (0.4) |
| Difference based variables (Flowering to maturity) |  |  |  |
| TMINMDIFF | Minimum temperature difference | -0.1 (0.9) | -4,-1.12 (0.5) |
| TMEANDIFF | Mean temperature difference | -0.3 (0.7) | -3.5, 1 (0.4) |
| TMAXDIFF | Maximum temperature difference | -0.4 (0.7) | -3, 1 (0.6) |
| DLMEANDIFF | Daylength difference | 0.2 (0.2) | -0.5, 0.7 (0.2) |
| Location based |  |  |  |
| Long | Longitude, deg | 23.1 (15.6) | -9.5, 38 (28.6) |
| Lat | Latitude, deg | -5.1 (11.4) | -20.5, 13.1 (21.7) |
| ALT | Altitude, m | 858 (470) | 148, 2160 (752) |

<sup>1</sup>SD= Standard deviation, <sup>2</sup> IQR= Interquartile range (difference between the 75<sup>th</sup> and 25<sup>th</sup> percentiles)

### S2. Modelling: Feature Selection

Beginning with one predictor at a time, stepwise forward regression was performed at (n+1) steps. No further improvement was noted after the seventh iteration, which included the features TMINMM, TMEANM, TMEANDIFF, DLMEANDIFF, ALT, lat, and long. To remove collinearity, models with one temperature -based predictor at the time along with DLMEANDIFF and geolocation (lat, long) were evaluated independently. Also, a model with ALT was considered. The best feature subset was:

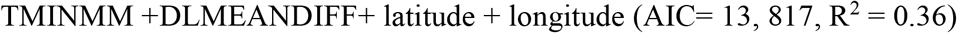

**Figure S4.**
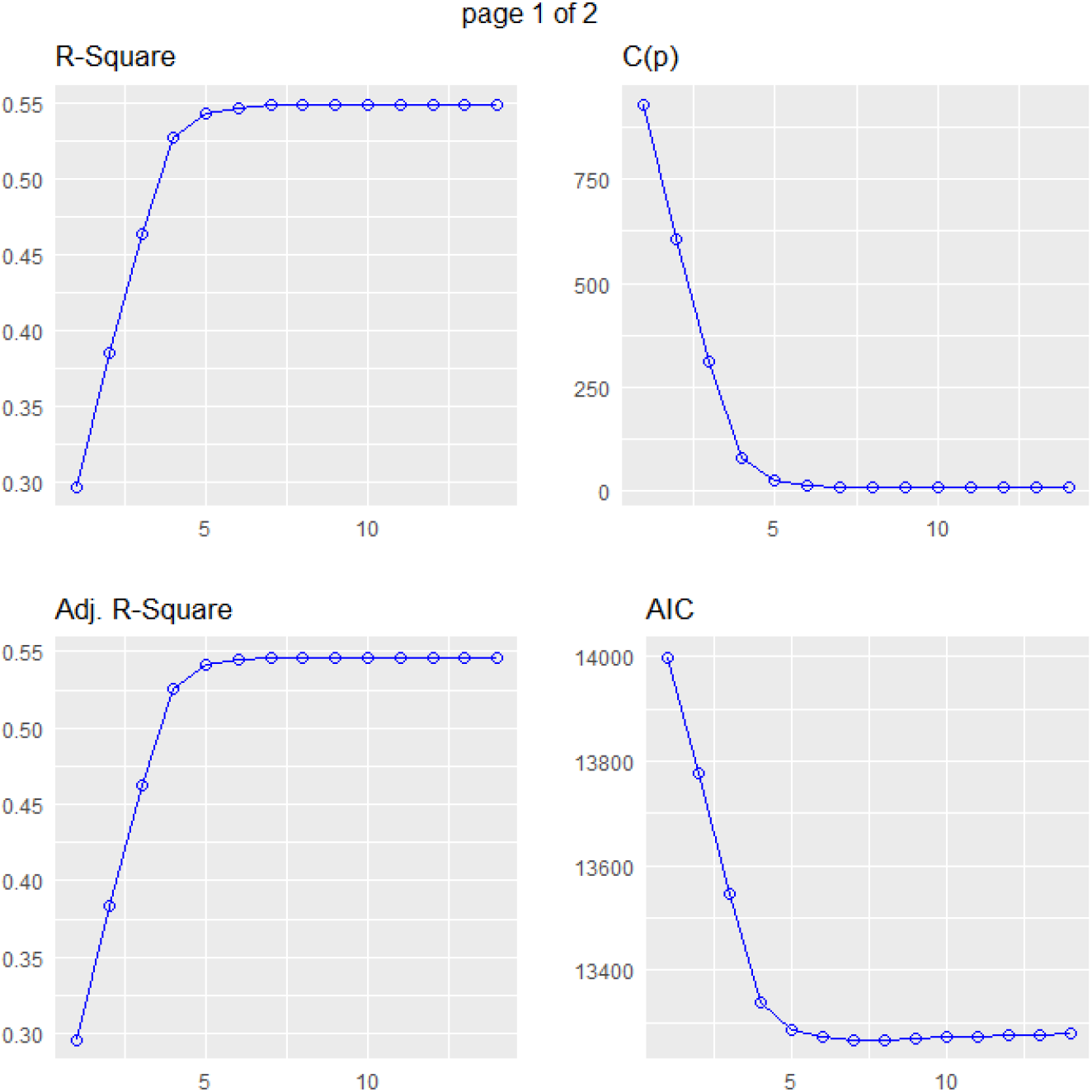
Stepwise forward selection for the best feature-set to predict maturity times using temperature-based variables. Model agreement improved no more after the seventh iteration. The best candidate features were TMINMM, TMEANM, TMEANDIFF, DLMEANDIFF, ALT, la t, and long.

**Table S2.** Evaluation and testing of GAM models used to predict Soybean maturity timing. Linear regression models (lm) included also for comparison.

| Model | 5-fold validation |  |  |  | All-train |  |
| --- | --- | --- | --- | --- | --- | --- |
|  | RMSE (days) |  | R<sup>2</sup>-adjusted |  |  |  |
|  | Training | Testing | Training | Testing | AIC | BIC |
| gam s(TMINM, 3) | 13.55 | 13.67 | 0.46 | 0.46 | 13542 | 13564 |
| gam s(TMINM, 6) | 13.17 | 13.31 | 0.49 | 0.48 | 16609 | 16646 |
| s(TMINM, 3) + s(lat, long, 4) | 13.17 | 13.35 | 0.48 | 0.48 | 16617 | 16657 |
| lm(TMINM + long) | -- | 16.24 | -- | 0.19 | 20057 | 20080 |
| gam s(TMINM,3)+s(long,4) | 13.14 | 13.32 | 0.49 | 0.48 | 16631 | 16670 |
| gam s(TMINM, 3) + long | 13.38 | 13.49 | 0.47 | 0.47 | 16669 | 16697 |
| gam s(TMINM, 6) + s(long, 6) | 11.95 | 12.30 | 0.58 | 0.56 | 16236 | 16304 |
| gam s(TMINM, 6) + s(long, 4) | 12.38 | 12.65 | 0.54 | 0.53 | 16370 | 16426 |
| gam s(TMINM, 6) + long | 12.54 | 12.73 | 0.53 | 0.53 | 16415 | 16460 |
| lm(TMINM+long+lat) | -- | 16.25 | -- | 0.18 | 20059 | 20087 |
| gam s(TMINM,6)+s(lat,long,4) | 12.43 | 12.67 | 0.54 | 0.53 | 16391 | 16497 |
| gam s(TMINM,6)+s(lat,long,6) | 12.41 | 12.67 | 0.54 | 0.53 | 16392 | 16457 |
| gam s(TMINM,6)+s(long,4)+s(lat,4) | 12.20 | 12.51 | 0.56 | 0.54 | 16322 | 16395 |
| gam s(TMINM,6)+s(long,4)+s(lat,6) | 11.80 | 12.08 | 0.59 | 0.57 | 16037 | 16122 |
| gam S(TMINM,6)+s(long,6)+s(lat,4) | 11.72 | 12.04 | 0.59 | 0.58 | 16191 | 16272 |
| gam s(TMINM,6)+s(long,6)+s(lat,6) | 11.65 | 12.01 | 0.60 | 0.58 | 15947 | 16043 |
| gam s(TMINM,6)+s(long,6)+lat | 11.95 | 12.23 | 0.57 | 0.56 | 16193 | 16266 |
| lm(TMINM+long+lat+DLMEANDIFF) | 14.90 | 15.48 | 0.34 | 0.32 | 13817 | 13850 |
| gam s(TMINM,4)+s(long,4)+s(lat,4)+DLMEANDIFF | 11.15 | 11.40 | 0.63 | 0.62 | 15944 | 16010 |
| gam s(TMINM,3)+s(long,6)+s(lat,6)+DLMEANDIFF | 10.76 | 11.10 | 0.65 | 0.64 | 15803 | 15587 |

Table S2. Evaluation and testing of GAM models used to predict Soybean maturity timing. Linear regression (lm) included also for comparison. (Continuation)
| Model | 5-fold validation |  |  |  | All-train |  |
| --- | --- | --- | --- | --- | --- | --- |
|  | RMSE (days) |  | R<sup>2</sup>-adjusted |  |  |  |
|  | Training | Testing | Training | Testing | AIC | BIC |
| lm(TMINM) | 15.43 | 15.79 | 0.29 | 0.30 | 14000 | 14016 |
| lm(TMINM+ DLMEANDIFF) | 14.96 | 15.48 | 0.32 | 0.33 | 13924 | 13945 |
| lm(TMINM + long) | -- | 16.24 | -- | 0.19 | 20057 | 20080 |
| gam s(TMINM, 3) + long | 13.38 | 13.49 | 0.47 | 0.47 | 16669 | 16697 |
| gam S(TMIN, 3) + DLMEANDIFF | 12.50 | 12.70 | 0.54 | 0.53 | 13280 | 13307 |
| gam S(TMIN, 3) + s( DLMEANDIFF,3) | 10.78 | 11.02 | 0.66 | 0.65 | 12793 | 12825 |
| gam s(TMINM, 6) + s(long, 6) | 11.95 | 12.30 | 0.58 | 0.56 | 16236 | 16304 |
| gam s(TMINM, 6) + s(long, 4) | 12.38 | 12.65 | 0.54 | 0.53 | 16370 | 16426 |
| lm(TMINM+long+lat) | -- | 16.25 | -- | 0.18 | 20059 | 20087 |
| gam s(TMINM,6)+s(lat,long,6) | 12.41 | 12.67 | 0.54 | 0.53 | 16392 | 16457 |
| gam s(TMINM,6)+s(long,4)+s(lat,6) | 11.80 | 12.08 | 0.59 | 0.57 | 16037 | 16122 |
| gam S(TMINM,6)+s(long,6)+s(lat,4) | 11.72 | 12.04 | 0.59 | 0.58 | 16191 | 16272 |
| gam s(TMINM,3)+s(long,6)+s(lat,6) | 11.65 | 12.01 | 0.60 | 0.58 | 15947 | 16043 |
| lm(TMINM+long+lat+DLMEANDIFF) | 14.90 | 15.48 | 0.34 | 0.32 | 13817 | 13850 |
| gam S(TMINM,3) + DLMEANDIFF | 12.50 | 12.70 | 0.54 | 0.53 | 13280 | 13307 |
| gam S(TMINM,3) + s(DLMEANDIFF,3) | 10.78 | 11.02 | 0.66 | 0.65 | 12793 | 12825 |
| gam s(TMINM,3)+s(long,6)+s(lat,6)+DLMEANDIFF | 10.76 | 11.10 | 0.65 | 0.64 | 15803 | 15587 |
| gam S(TMINM,3) + S(DLMEANDIFF,3 )+lat + long | 10.05 | 10.3 | 0.70 | 0.69 | 12576 | 12620 |
| gam S(TMIN,3)+s(DLMEANDIFF,3)+s(lat,long,4) | 9.99 | 10.35 | 0.70 | 0.69 | 12564 | 12613 |

The indicator s is the basis function with k-break points or knots. e.g., s(TMINM, 3) is the response of maturity to TMINM with 3 knots

### S3. Time to maturity Cultivar x Environment Interactions

**Figure S5.**
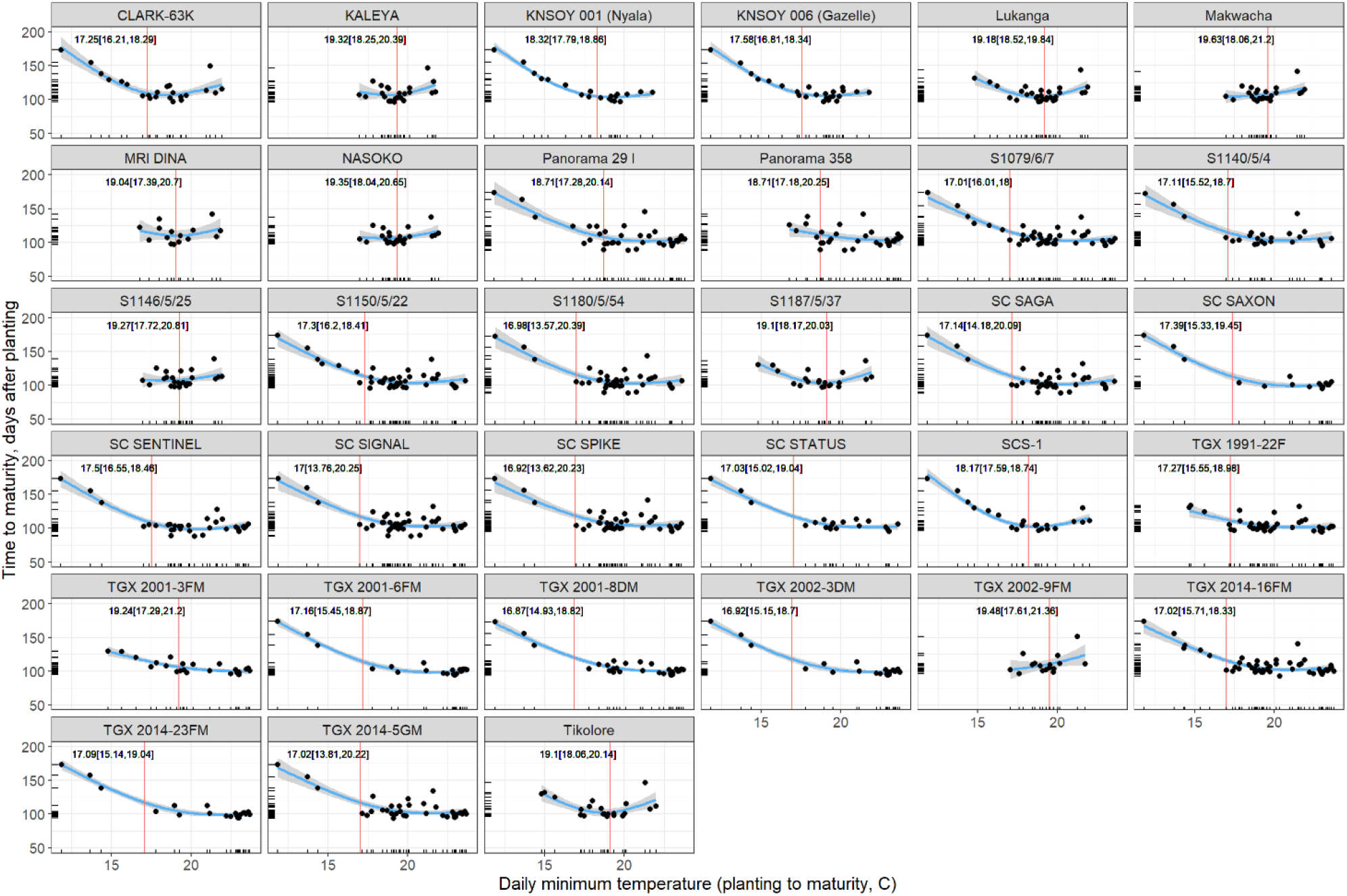
GAM smoothed response of soybean maturity time to minimum temperature. Each dot represents a testing site where the cultivar was tested. Red parallel lines represent a change in the direction of the response, approximated using segmented regression. 33 cultivars showed a significant relationship.

**Figure S6.**
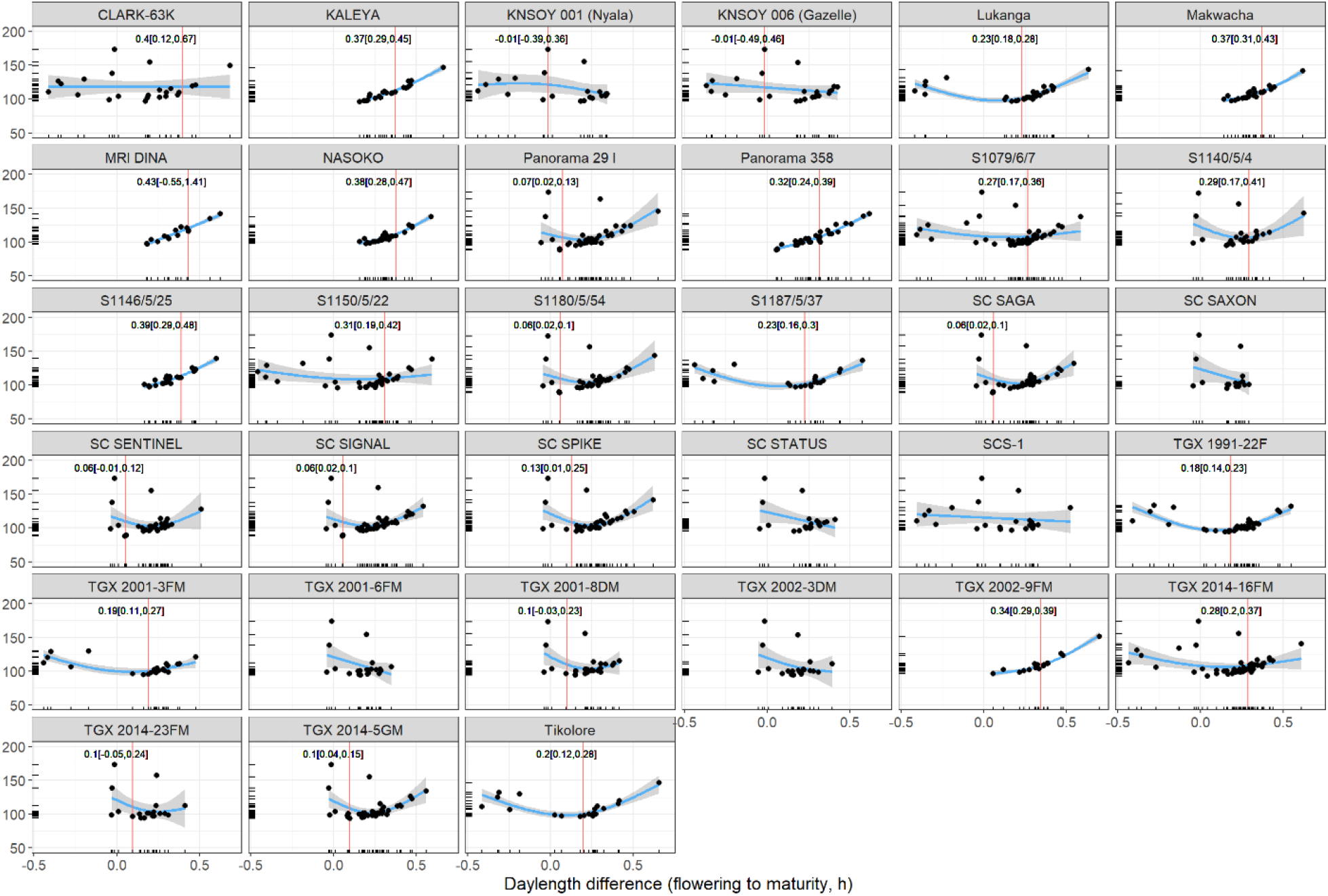
GAM smoothed response of soybean maturity time to post-flowering daylength. Each dot represents a testing site where the cultivar was tested. Red parallel lines represent a change in the direction of the response, approximated using segmented regression. 5 out of 33 cultivars failed to display a significant change in response.

